# Precise regulation of gene expression through transcriptional repression is essential for *Plasmodium* asexual blood stage development

**DOI:** 10.1101/2025.04.30.651575

**Authors:** Tsubasa Nishi, Izumi Kaneko, Shiroh Iwanaga, Masao Yuda

## Abstract

Malaria is caused by the proliferation of *Plasmodium* parasites in the vertebrate host blood stream through repeated cycles of asexual multiplication inside erythrocytes. During these cycles, parasites dynamically change their transcriptome at each developmental step to express genes exactly when required; however, the mechanisms regulating these transcriptomic changes remain unclear. In this study, we revealed that the AP2-family transcription factor PbAP2-TR is essential for the asexual blood stage development of the rodent malaria parasite *Plasmodium berghei*, as a transcriptional repressor. Conditional knockout of *pbap2-tr* caused developmental arrest at the trophozoite stage, *i.e.*, the cell growth phase of asexual blood stage development. Chromatin immunoprecipitation followed by high-throughput sequencing showed that PbAP2-TR binds to two different DNA motifs and targets genes that are downregulated towards late trophozoites, including ribosome biogenesis-related genes and host-modifying exported protein genes. The expression of these target genes was upregulated in *pbap2-tr*-knockout parasites, and introduction of mutations into the binding motifs increased the promoter activity of the target genes. These results indicate that PbAP2-TR establishes precise transcription peak patterns of its target genes by repressing their transcription during the trophozoite, thereby being essential for asexual blood stage development. Our data also suggest that PbAP2-TR induces transcriptional repression by recruiting a putative chromatin remodeler, PbMORC, as a co-factor.

## Introduction

*Plasmodium* is the causative agent of malaria, which is one of the most serious infectious diseases worldwide (1). The parasite proliferates in host blood through repeated cycles of red blood cell (RBC) invasion by a specific invasive stage called the merozoite and asexual replication inside host RBCs (2–4). Intraerythrocytic asexual development can be divided into three stages: ring, trophozoite, and schizont (5). The ring stage, which is the first developmental stage formed after merozoite invasion, is characterized by a biconcave disc shape (6). At the ring stage, the expression of exported proteins is upregulated to modify the host cell into a parasite-favorable environment (7). The next developmental stage, trophozoite, grows rapidly with an elevated uptake of host-derived nutrients, including hemoglobin, which is a source of amino acids (8, 9). For rapid cell growth, gene transcription becomes considerably more active during the trophozoite stage than during the ring stage (10). The parasite then develops into the schizont stage, which undergoes a unique mode of cell division called schizogony (11, 12). In schizonts, several rounds of asynchronous nuclear division occur while merozoite components, such as the rhoptry and inner membrane complex (IMC), are generated (13). Finally, multiple progeny merozoites are formed through segmentation simultaneously with a final round of nuclear division.

As the asexual blood stage development proceeds, the parasite dynamically alters gene expression. Time-course transcriptome analyses using microarray and steady-state RNA sequencing (RNA-seq) have shown that the transcript levels of genes that have a common functional role show similar peak patterns in the course of the intraerythrocytic developmental cycle (IDC) (14–16). Such temporal changes in gene transcription largely appear as the cascade of transcriptional activation to express genes just when they are required, *i.e.*, “just-in-time” transcription. Concordant with the transcription peak patterns, nucleosome-free or nucleosome-depleted regions, which indicate active transcription sites, are also periodically established (17). Furthermore, a recent study indicated the essentiality of chromatin remodelers for the “just-in-time” regulation of gene expression during the asexual blood stage development (18). These suggest that regulation at the transcriptional level is important for IDC progression; thus, investigating the mechanisms that underlie this periodic transcriptional regulation is very important to obtain a deep understanding of parasite asexual proliferation in the host blood.

*Plasmodium* spp. have a limited number of sequence-specific transcription factors encoded in their genome. These transcription factors are mainly AP2 family proteins, which are homologues of the plant APETALA2/Ethylene Response Factor (AP2/ERF) and contain one to three DNA-binding domains called AP2 domains (19). In previous studies, several stage-specific transcription factors have been explored, and extensive progress has been particularly made regarding the development of invasive and sexual stages (20–30). At these stages, transcriptional activation of stage-specific genes is controlled by a few sequence-specific transcription factors and their corresponding binding motifs. For the asexual blood stage development, two transcriptional activators have been reported, AP2-I and SIP2 (23, 28). In *P. falciparum*, PfAP2-I is expressed from trophozoites to early schizonts, and targets invasion-related genes during the schizont stage. In *P. berghei*, PbSIP2 functions as a master regulator essential for merozoite formation, comprehensively activating genes related to merozoite formation, such as genes related to the rhoptry, microneme, and IMC. Meanwhile, transcription factors that control transcriptomic changes during the other stages of the IDC, *i.e.* ring and trophozoite stages, have not been identified, and the mechanism regulating the transcription peak patterns for each gene remains largely unknown.

To advance our knowledge of transcriptional regulation during the IDC, we focused on AP2 transcription factor genes that could not be knocked out during the IDC of *P. berghei*. Here, we demonstrate that the AP2 family transcription factor, PbAP2-TR (a trophozoite repressor), functions as a transcriptional repressor essential for trophozoite development. Our data reveal that PbAP2-TR represses the transcription of its target genes during trophozoite development and controls their transcriptional profiles.

## Results

### PbAP2-TR is an AP2 transcription factor expressed in trophozoites

PbAP2-TR is an AP2-family transcription factor that possesses three conserved AP2 domains (Fig 1A). The first (position 182–235) and second (position 279–325) AP2 domains are located in tandem near the N-terminus, and the third (position 1474–1522) domain is located towards the C-terminus. When the amino acid (AA) sequences of the tandem AP2 domains were aligned among PbAP2-TR orthologs in *Plasmodium* species, both the AP2 domains and the linker between them were found to be highly conserved (96% and 87% for the first and second AP2 domains, respectively; 81% for the linker) (Fig 1B). For the third AP2 domain of PbAP2-TR, the AA sequences were completely conserved within *Plasmodium* (Fig 1C). In addition to these AP2 domains, PbAP2-TR contains a conserved AP2-coincident domain mostly at the C-terminus (ACDC) and a putative nuclear localization signal (NLS) at position 1139–1151 (Fig 1A).

**Figure 1.**
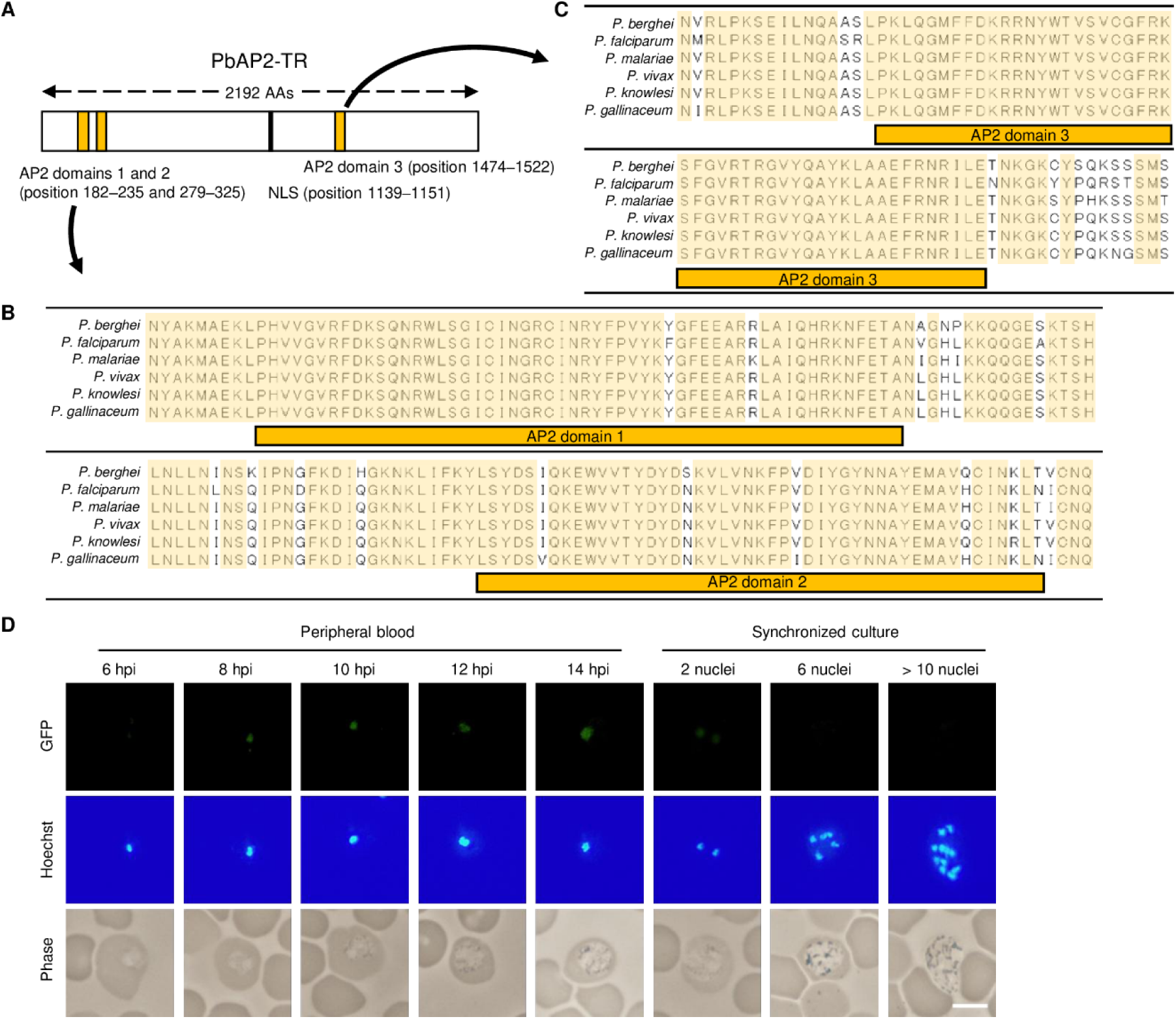
Structural features of PbAP2-TR and its expression pattern during the asexual blood stage development. (A) Schematic illustration of PbAP2-TR. The regions of an AP2 domain are indicated by yellow. The nuclear localization signal (NLS) was predicted using cNLS Mapper (http://nls-mapper.iab.keio.ac.jp/cgi-bin/NLS_Mapper_form.cgi) and is indicated with a black bar. (B) Alignment of amino acid sequences for the two tandem AP2 domains of PbAP2-TR orthologs in *Plasmodium* using the ClustalW program in Mega X (*P*. *berghei*, PBANKA_0909600; *P*. *falciparum*, PF3D7_1139300; *P. malariae*, PmUG01_09048500; *P. vivax*, PVP01_0940100; *P*. *knowlesi*, PKNH_0937300; *P. gallinaceum*, PGAL8A_00366800). Amino acids conserved in all orthologs are highlighted by yellow. The regions of an AP2 domain are indicated by yellow boxes. (C) Alignment of amino acid sequences for the third AP2 domain of PbAP2-TR orthologs in *Plasmodium*. (D) Expression of PbAP2-TR in the PbAP2-TR::GFP during asexual blood stage development. Time-course analysis was performed using the peripheral blood of infected mice. Schizonts were observed in the cultures. Nuclei were stained with Hoechst 33342. Scale bar = 5 μm.

We first attempted to generate *pbap2-tr*-knockout parasites using conventional homologous recombination (31) but failed to obtain mutant parasites. This was consistent with previous knockout screening studies on rodent malaria parasites (32–34). These results indicate that *pbap2-tr* is essential for asexual blood stage development. To evaluate at which cell-stage PbAP2-TR functions during intraerythrocytic development, we assessed the PbAP2-TR expression pattern. We generated a parasite line expressing GFP-fused PbAP2-TR (PbAP2-TR::GFP, Fig S1) using the Cas9-expressing parasite, PbCas9 (35), and performed a time-course fluorescent analysis. The cell cycle was synchronized to mature schizonts by incubating PbAP2-TR::GFP-infected blood in culture for 16 h, and the cultured schizonts were then intravenously injected into mice. A nuclear-localized GFP signal was first observed in early trophozoites at 8 h post-injection (hpi) (Fig 1D). The signal continued to be observed for the late trophozoites in the peripheral blood (Fig 1D). As later asexual blood stages are rarely observed in peripheral blood owing to sequestration, we further performed fluorescence analysis on cultured PbAP2-TR::GFP. In the culture, mononuclear parasites showed nuclear-localized GFP signals, as observed in peripheral blood; however, the signal was considerably weaker in schizonts with two nuclei. Furthermore, later schizonts showed no GFP signal (Fig 1D). These results indicate that PbAP2-TR is a trophozoite-specific transcription factor.

### Disruption of *pbap2-tr* results in developmental arrest at the trophozoite stage

As gene disruption of *pbap2-tr* was unsuccessful, we performed a conditional knockout of *pbap2-tr* to assess the role of PbAP2-TR. We employed a previously reported dimerizable Cre (DiCre) system (36, 37) using a parasite line that constitutively expressed Cas9 and DiCre (PbCas9^DiCre^) (28). Two loxP sequences were each introduced to the 5′ and 3′ sides of *pbap2-tr*, arranged in the same direction (*pbap2-tr*-DiCre, Fig S2A), for removing the *pbap2-tr* locus in a rapamycin-dependent manner (Fig 2A). For the conditional knockout, whole blood was harvested from mice infected with *pbap2-tr*-DiCre and split into two cultures (Fig 2B). At the beginning of the culture, rapamycin was added to one culture (final concentration, 30 nM) (*pbap2-tr*-DiCre^Rapa+^), and dimethyl sulfoxide (DMSO) was added to the other (*pbap2-tr*-DiCre^Rapa-^) as a control (Fig 2B). After 16 h of culture, genotyping PCR confirmed that the *pbap2-tr* locus was almost completely excised from the genome of *pbap2-tr*-DiCre^Rapa+^, with only a subtle signal for the wild-type (WT) *pbap2-tr* locus (Fig 2C). To assess the effect of *pbap2-tr*-disruption, mature schizonts of *pbap2-tr*-DiCre^Rapa+^ and *pbap2-tr*-DiCre^Rapa-^ were inoculated into mice (Fig 2B). The parasites were then cultured at 10 hpi, and parasite development was assessed by Giemsa-staining every four hours (Fig 2B). In the cultures, *pbap2-tr*-DiCre^Rapa-^ and *pbap2-tr*-DiCre^Rapa+^ showed similar ring-to-trophozoite ratios until 18 hpi (Fig 2D). In addition, the average size of trophozoites at 18 hpi was comparable between *pbap2-tr*-DiCre^Rapa-^ and *pbap2-tr*-DiCre^Rapa+^ with a diameter of approximately 4.4 μm (Fig 2E). However, at 22 hpi, the schizont ratio reached 20% in *pbap2-tr*-DiCre^Rapa-^, whereas most of the *pbap2-tr*-DiCre^Rapa+^ parasites remained as mononuclear cells (schizont ratio: 0.6%) (Fig 2D). Furthermore, the ratio of schizonts in *pbap2-tr*-DiCre^Rapa+^ was still less than 3% at 26 hpi, whereas that in *pbap2-tr*-DiCre^Rapa-^ was higher than 60% (Fig 2D). This suggests that PbAP2-TR functions during trophozoite development before the beginning of nuclear division.

**Figure 2.**
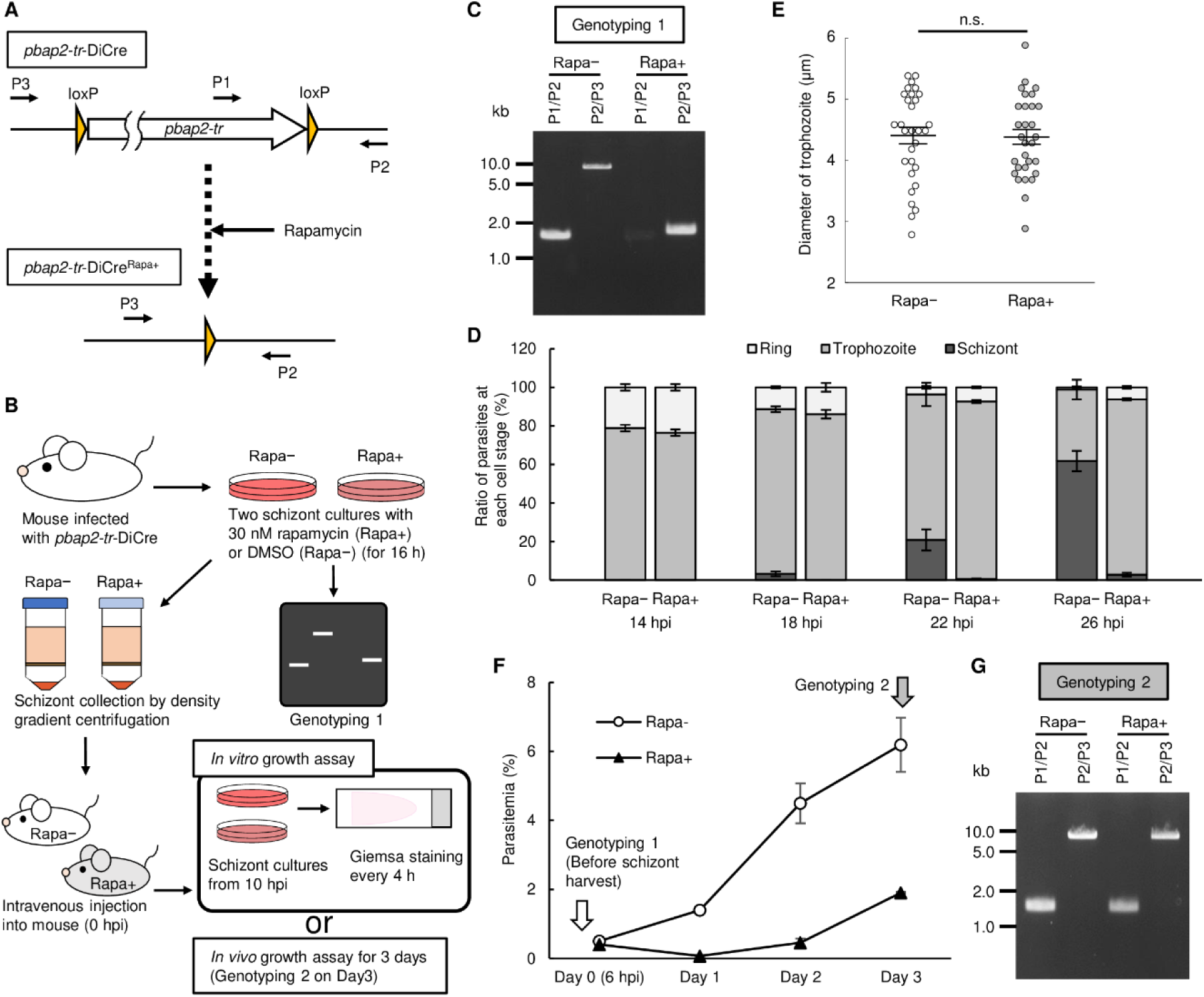
Conditional knockout of *pbap2-tr* using the DiCre system. (A) Schematic illustration of the *pbap2-tr* locus of *pbap2-tr*-DiCre and the DiCre-mediated recombination induced by rapamycin treatment. Arrows P1–P3 indicate the locations of primers designed for genotyping PCR. (B) Outline of the conditional knockout experiments of *pbap2-tr* using the DiCre system. First, whole blood from mice infected with *pbap2-tr*-DiCre was split into two cultures: one with 30 nM rapamycin (Rapa+) and the other with DMSO as a control (Rapa−). After 16 h of culture, schizonts were harvested by density gradient centrifugation and intravenously injected into mice (0 hours post-injection (hpi)). Parasite growth was then assessed *in vitro* or *in vivo*. For assessing *in vitro*, whole blood was harvested from the infected mice at 10 hpi and cultured again. Parasite development was then assessed by Giemsa-staining every 4 hours. For assessing *in vivo*, parasitemia was assessed daily using the peripheral blood from the mice infected with rapamycin- or DMSO-treated *pbap2-tr*-DiCre. (C) Representative gel image of the genotyping PCR analysis for *pbap2-tr*-DiCre^Rapa−^ and *pbap2-tr*-DiCre^Rapa+^ performed at 16 h after starting the culture (Genotyping 1). The primers used are illustrated in (A). (D) Ratio of ring, trophozoite, and schizont stage parasites at each time point of *pbap2-tr*-DiCre^Rapa−^ and *pbap2-tr*-DiCre^Rapa+^ cultures. Parasite developmental stages were determined by Giemsa staining. Error bars indicate the standard error of the mean value from three biologically independent experiments. (E) Size of trophozoites for *pbap2-tr*-DiCre^Rapa−^ and *pbap2-tr*-DiCre^Rapa+^ at 18 hpi. The longest diameter (μm) for each trophozoite was measured on Giemsa-stained blood smears. Lines indicate the mean values and the standard error (n = 30). (n.s.: not significant, the *p*-value calculated using two-tailed Student’s t-test was higher than 0.05.) (F) Parasite growth of *pbap2-tr*-DiCre^Rapa−^ and *pbap2-tr*-DiCre^Rapa+^ *in vivo*. Parasitemia was assessed by Giemsa staining. Error bars indicate the standard error of the mean value from three biologically independent experiments. Time points for Genotyping 1 and 2 are indicated by arrows. (G) Representative gel image of the genotyping PCR analysis for *pbap2-tr*-DiCre^Rapa−^ and *pbap2-tr*-DiCre^Rapa+^ performed on day 3 (Genotyping 2).

To further confirm the essentiality of *pbap2-tr* in asexual blood stage development, we assessed the parasite growth of *pbap2-tr*-DiCre *in vivo*. After inducing rapamycin-dependent knockout and inoculating the schizonts of *pbap2-tr*-DiCre^Rapa-^ and *pbap2-tr*-DiCre^Rapa+^ into mice as, parasitemia was determined through daily Giemsa staining. Parasite infectivity immediately after inoculation was comparable between *pbap2-tr*-DiCre^Rapa-^ and *pbap2-tr*-DiCre^Rapa+^ as their parasitemia was not significantly different on day 0 (6 hpi), suggesting that *pbap2-tr*-disruption did not affect the ability of schizonts/merozoites to infect host RBCs (Fig 2F). On day 1, the parasitemia of *pbap2-tr*-DiCre^Rapa+^ decreased from that on day 0 and was significantly lower than that of *pbap2-tr*-DiCre^Rapa-^ (Fig 2F). In the following days, parasitemia of *pbap2-tr*-DiCre^Rapa+^ increased with a growth rate comparable to that of *pbap2-tr*-DiCre^Rapa-^ (Fig 2F). However, genotyping on day 3 did not detect *pbap2-tr*-disrupted parasites but only parasites with the WT *pbap2-tr* locus (Fig 2G). This result confirms that *pbap2-tr* plays an essential role in parasite development during the IDC.

Previously, Shang *et al*. reported that in *P. falciparum*, disruption of the *pbap2-tr* ortholog, named *pfap2-g5*, resulted in upregulation of PfAP2-G and increased the rate of differentiation into gametocytes, consequently causing a significant reduction in the parasite asexual growth rate (38). Thus, they concluded that PfAP2-G5 is important for suppressing commitment to the gametocyte fate. To evaluate whether PbAP2-TR plays a similar role, we disrupted *pbap2-g* in *pbap2-tr*-DiCre using the CRISPR/Cas9 system [*pbap2-tr*-DiCre*^pbap2-g^*^(-)^, Fig S2B]. DiCre-mediated disruption of *pbap2-tr* was induced in cultures (*pbap2-tr*-DiCre*^pbap2-g^*^(-)_Rapa+^ and *pbap2-tr*-DiCre*^ap2-g^*^(-)_Rapa-^) (Genotyping 1, Fig S2C left), and after inoculating *pbap2-tr*-DiCre*^pbap2-g^*^(-)_Rapa-^ and *pbap2-tr*-DiCre*^ap2-g^*^(-)_Rapa+^ into mice, parasitemia was assessed daily. The result was similar to that of the phenotype analysis for *pbap2-tr*-DiCre; in *pbap2-tr*-DiCre*^ap2-g^*^(-)_Rapa+^, parasitemia decreased from day 0 to day 1 and later increased (Fig S2D), but the day-3 parasites possessed the WT *pbap2-tr* locus (Genotyping 2, Fig S2C right). This result indicates that in *P. berghei*, PbAP2-TR is essential for asexual blood stage development and that repression of *ap2-g* transcription is not a common function of PbAP2-TR orthologs.

### PbAP2-TR recognizes two DNA motifs using different AP2 domains

To investigate the target genes regulated by PbAP2-TR, we performed chromatin immunoprecipitation followed by high-throughput sequencing (ChIP-seq) of PbAP2-TR at the trophozoite stage. To specifically assess the role of PbAP2-TR in asexual development, we disrupted *pbap2-g* in PbAP2-TR::GFP using the CRISPR/Cas9 system [PbAP2-TR::GFP*^pbap2-g^*^(-)^, Fig S3]. ChIP-seq analyses were performed in duplicate using PbAP2-TR::GFP*^pbap2-g^*^(-)^ at 12 hpi. Experiments 1 and 2 detected 1086 and 1259 peaks, respectively, and 1017 peaks overlapped (93% of the peaks in experiment 1) (Fig 3A, Table S1A, and S1B). From the common peaks, we first explored the putative binding motifs of PbAP2-TR by DNA motif enrichment analysis using Fisher’s exact test. The analysis revealed GTTGTA as the most enriched motif (*p*-value = 5.2 × 10^−72^) and several motifs containing GTTGT in the 20 most enriched motifs (Table S1C). Furthermore, the analysis detected the enrichment of motifs containing GTTGC. Thus, the most enriched motif was considered as GTTGY (Fig 3B). Among the 1017 ChIP-seq peaks, 872 peaks contained the GTTGY motif within 300 bp of the summit, and most were found within 100 bp (Fig 3C). In addition to GTTGY, motifs containing GTTCT were highly enriched (Fig 3D). This motif was found in 514 peaks, mostly close to the summit (Fig 3E). Hereafter, the GTTGY and GTTCT motifs are referred to as Motif 1 and 2, respectively. In total, 90% of the peaks (934 peaks) contained either or both Motif 1 and 2.

**Figure 3.**
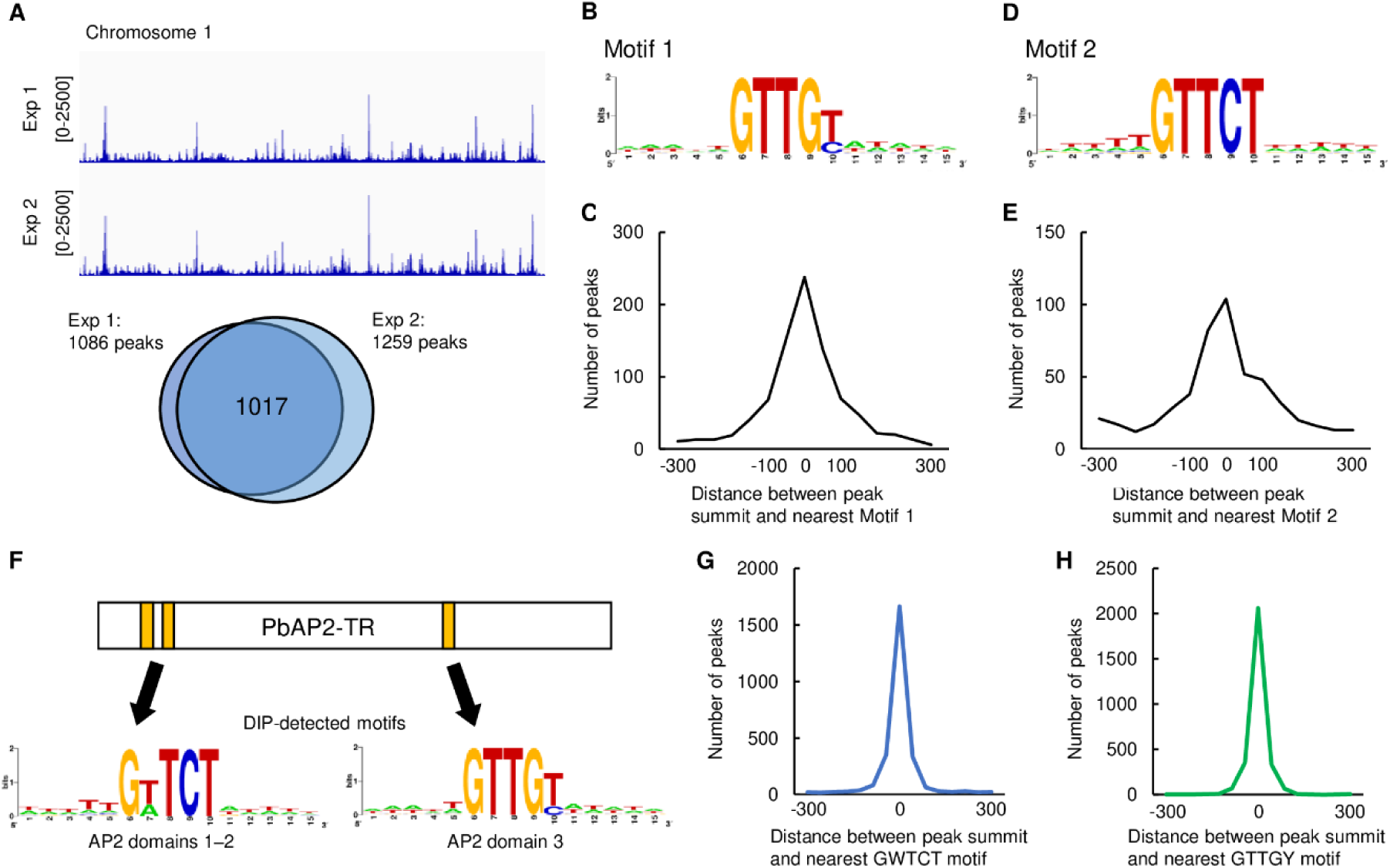
DNA binding properties of PbAP2-TR assessed by chromatin immunoprecipitation followed by high-throughput sequencing (ChIP-seq) and DNA immunoprecipitation followed by high-throughput sequencing (DIP-seq) analyses. (A) Integrative Genomics Viewer images for the PbAP2-TR ChIP-seq experiments 1 and 2 on chromosome 1. Histograms show the row read coverage of ChIP data at each base. Scales are indicated in square brackets. A Venn diagram at the bottom shows the number of overlapping peaks between experiments 1 and 2. (B) The motif most enriched within 50 bp from the summits of the PbAP2-TR ChIP-seq peaks (Motif 1). The logo was generated using WebLogo (https://weblogo.berkeley.edu/logo.cgi). (C) Distance between the peak summits and the nearest Motif 1. (D) The second most enriched motif for PbAP2-TR ChIP-seq peaks (Motif 2). (E) Distance between the peak summits and the nearest Motif 2. (F) Motifs enriched in the DIP-seq peak regions for AP2 domains 1–2 (left) and domain 3 (right). The schematic illustration of PbAP2-TR is shown at the top. The logos were generated using WebLogo. (G) Distance between DIP-seq peak summits for AP2 domains 1–2 and the nearest GWTCT (W = A or T) motifs. (H) Distance between the DIP-seq peak summits for AP2 domain 3 and the nearest GTTGY motifs.

To further evaluate the DNA-binding properties of PbAP2-TR, we performed DNA immunoprecipitation followed by high-throughput sequencing (DIP-seq) analysis using recombinant PbAP2-TR AP2 domains fused with maltose-binding protein (MBP) and fragmented *P. berghei* genomic DNA. First, DIP-seq was performed on two N-terminal tandem AP2 domains. Because these two AP2 domains likely function together owing to their proximity and the conserved amino acids between them (Fig 1B), we generated a recombinant protein containing both AP2 domains. In this analysis, 3038 peaks were detected throughout the genome (Table S2A). Motif enrichment analysis of these DIP peaks detected GTTCT and GATCT as the two most enriched motifs (Fig 3F and Table S2B). Among the 3038 DIP peaks, 2502 peaks (82.3%) contained GWTCT (W = A or T) within 100 bp of their summit (Fig 3G), suggesting that GWTCT, which is analogous to Motif 2, is the binding motif of the N-terminal tandem AP2 domains of PbAP2-TR. Next, DIP-seq was performed on the third AP2 domain of PbAP2-TR. In this analysis, 2980 peaks were detected (Table S2C), and the most enriched motif in these peak regions was GTTGT (Table S2D). The following enriched motifs were one-base-shifted motifs of GTTGT (TGTTG, AGTTG, and TTGTA) and GTTGC (Table S2D). Thus, the enriched motif was identical to Motif 1 (GTTGY) (Fig 3F). Motif 1 was contained within 100 bp from the summit of 2888 peaks (97%) (Fig 3H). Collectively, these results suggest that PbAP2-TR binds to Motif 1 and 2 each by different AP2 domains.

### PbAP2-TR targets are downregulated during early to late trophozoite development

Based on ChIP-seq data, we identified 619 target genes of PbAP2-TR, which were defined as genes with a ChIP-seq peak within 1200 bp of the start codon (Table S3). To examine the role of PbAP2-TR in transcriptional regulation, we investigated the changes in the transcript levels of these target genes from the beginning (8 hpi) to the end (16 hpi) of PbAP2-TR expression using high-throughput RNA sequencing (RNA-seq). To obtain asexual transcriptomic data, RNA-seq analyses were conducted using gametocyte-absent mutant parasites, *i.e.*, the *ap2-g*-knockout line. Among the genes significantly downregulated at 16 hpi compared with those at 8 hpi (562 genes in total, log_2_(fold change) > −1, *p*-value adjusted for multiple testing with the Benjamini-Hochberg procedure (*p*-value^adj^) < 0.05), 170 genes were PbAP2-TR targets whereas only 44 targets were included in the significantly upregulated genes (544 genes in total, log_2_(fold change) < 1, *p*-value^adj^ < 0.05) (Fig 4A and Table S4). Overall, the average log_2_(fold change) value of the PbAP2-TR target genes was significantly lower than that of the other genes (*p*-value = 2.6 × 10^−34^ by two-tailed Student’s t-test). In addition, in the upstream region (200–1000 bp from the start codon) of the downregulated genes, GTTGT (Motif 1) was the most significantly enriched with a *p*-value of 9.6 × 10^−34^ by Fisher’s exact test (Fig 4B). The other binding motifs, GTTGC (Motif 1) and GTTCT (Motif 2), were also enriched with *p*-values of 1.2 × 10^−9^ and 5.2 × 10^−2^, respectively (Fig 4B). These results suggest that PbAP2-TR is the major transcriptional repressor responsible for gene downregulation during early to late trophozoite development.

**Figure 4.**
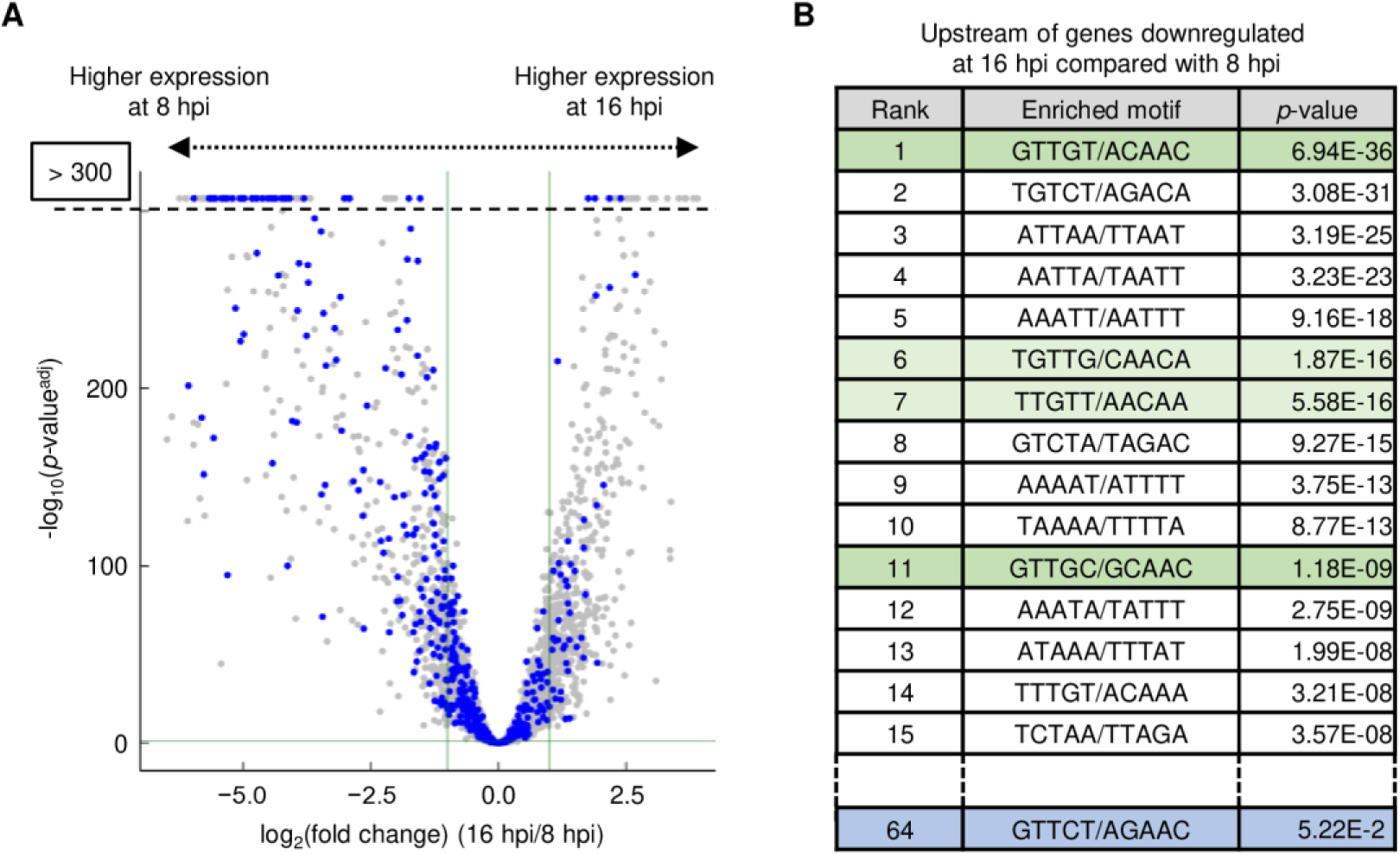
Transcriptional changes in PbAP2-TR targets during the PbAP2-TR expression period. (A) Volcano plot showing the differential expression of genes from 8 to 16 hpi. Blue dots represent the PbAP2-TR targets. The horizontal line indicates a *p*-value of 0.05 and the two vertical lines indicate the log_2_(fold change) of 1 and −1. Genes located higher than the horizontal dotted line have a *p*-value less than 10^-300^. (B) DNA motifs enriched in the upstream regions (200–1000 bp from the start codons) of genes downregulated from 8 to 16 hpi. Motif 1 (GTTGY) and Motif 2 (GTTCT) are indicated in green and blue, respectively. Motifs partially overlapped with Motif 1 are indicated in light green.

### PbAP2-TR functions as a transcriptional repressor

To verify that PbAP2-TR is a transcriptional repressor, we performed a differential expression analysis between *pbap2-tr*-DiCre^Rapa-^ and *pbap2-tr*-DiCre^Rapa+^ at 12 hpi. The analysis revealed that 70 and 111 genes were significantly upregulated and downregulated, respectively, in *pbap2-tr*-DiCre^Rapa+^ compared with those in *pbap2-tr*-DiCre^Rapa-^ (Fig 5A and Table S5). Among the upregulated genes, 41 PbAP2-TR target genes were detected, showing significant enrichment with a *p*-value of 1.1 × 10^−18^ by Fisher’s exact test (Fig 5A). In contrast, the downregulated genes included only nine target genes (Fig 5A). Furthermore, the average log_2_(fold change) value for the PbAP2-TR target genes was significantly higher than that for the other genes with a *p*-value of 4.8 × 10^−65^ by two-tailed Student’s t-test (Fig 5B). Motif enrichment analysis showed enrichment of both Motif 1 and 2 in the upstream sequences (200–1000 bp from the start codon) of the upregulated genes compared with those of the other genes (Fig 5C). These results confirm that PbAP2-TR functions as a transcriptional repressor during trophozoite development.

**Figure 5.**
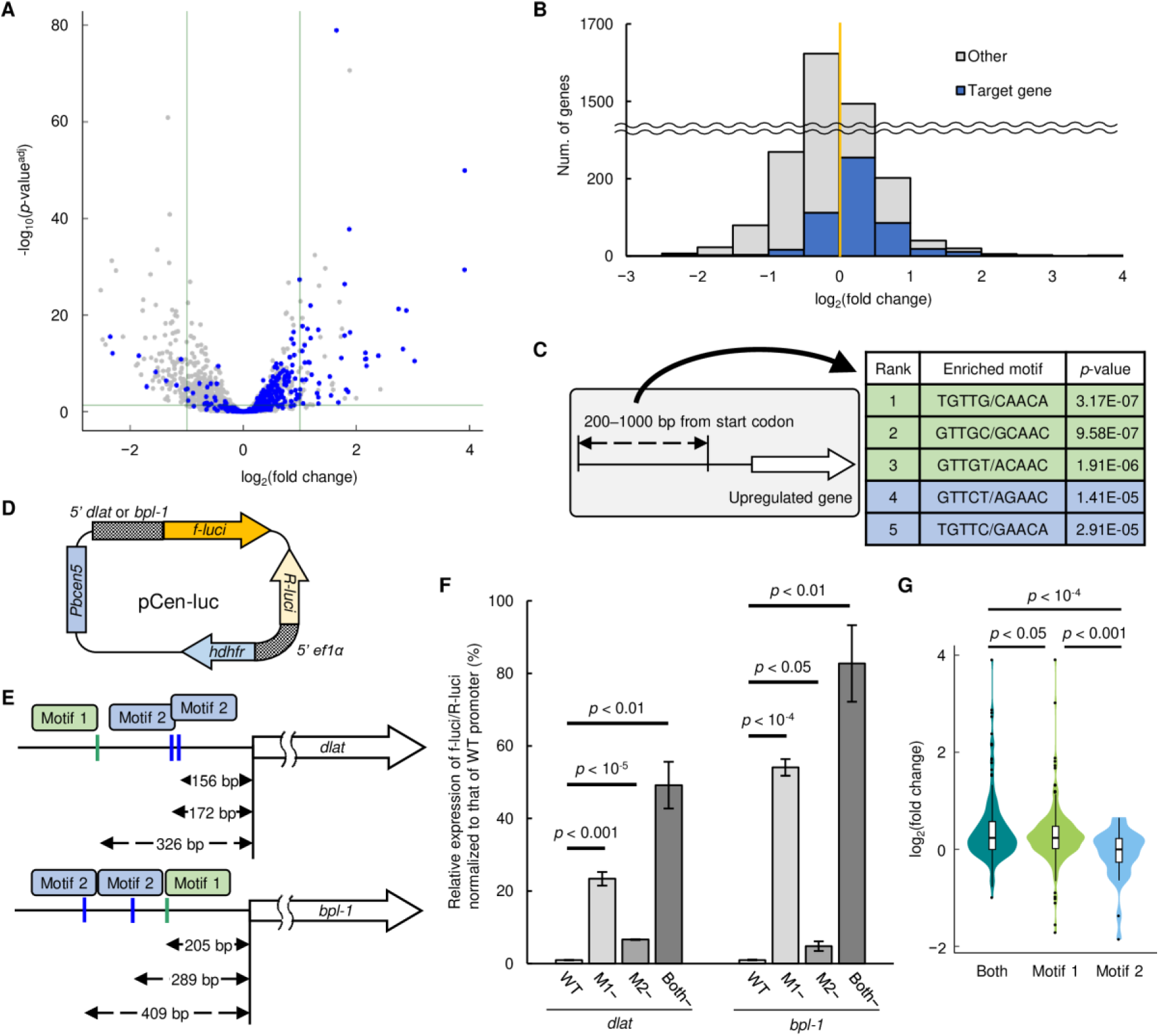
Function of PbAP2-TR as a transcriptional repressor. (A) Volcano plot showing the differential expression of genes between *pbap2-tr*-DiCre^Rapa−^ and *pbap2-tr*-DiCre^Rapa+^ at 12 hours post-injection. Blue dots represent the target genes of PbAP2-TR. The horizontal line indicates a *p*-value of 0.05 and the two vertical lines indicate the log_2_(fold change) of 1 and −1. (B) Histogram showing the number of genes against the log_2_(fold change) values. The number of PbAP2-TR targets for each bin is indicated in blue. The yellow line indicates a log_2_(fold change) of 0. (C) Five most enriched DNA motifs in the upstream regions (200–1000 bp from start codons) of genes upregulated in *pbap2-tr*-DiCre^Rapa+^. Motifs related to Motif 1 (GTTGY) and Motif 2 (GTTCT) are indicated in green and blue, respectively. (D) Schematic illustration of the pCen-luc plasmid (*f-luci*, firefly luciferase gene; *R-luci*, *Renilla* luciferase gene; *hdhfr*, human dihydrofolate reductase gene; *5′ ef1α*, bi-directional promoter of *P. berghei* elongation factor 1-alpha genes; *Pbcen5*, the centromere of *P. berghei* chromosome 5). (E) Schematic illustrations of the *dlat* and *bpl-1* loci. Locations of Motif 1 (GTTGY) and 2 (GTTCT) are indicated in green and blue, respectively, with distances from the start codon of each gene. (F) Dual-luciferase reporter assays for the promoter activities of *dlat* and *bpl-1*. The relative luminescence intensity (f-luci/R-luci) for each reporter was normalized to that of the corresponding wild-type reporter (WT). M1− and M2− are reporters with mutations in Motif 1 and Motif 2, respectively. Both− is a reporter with mutations in both Motifs 1 and 2. Error bars indicate the standard error of the mean value from three biologically independent experiments. The *p*-values were calculated using a two-tailed Student’s t-test (n.s.: not significant, *p*-value > 0.05). (G) Violin plot showing the distribution of log_2_(fold change) values for three groups of PbAP2-TR targets: those with a ChIP-seq peak containing both Motif 1 and 2, only Motif 1, and only Motif 2. The *p*-values were calculated using a two-tailed Student’s t-test. The corresponding box plots are shown inside the violin plots, and the dots indicate outliers.

### Binding motifs of PbAP2-TR function as *cis*-regulatory repressing elements

To evaluate the functions of Motif 1 and 2 as *cis*-regulatory elements, we performed dual luciferase assays using a centromere plasmid containing firefly and *Renilla* luciferase genes (pCen-luc, Fig 5D). Expression of the *Renilla* luciferase (R-luci) gene in pCen-luc was controlled by the *P. berghei* elongation factor 1-alpha promoter as an internal control, and the target gene promoters were inserted upstream of the firefly luciferase (f-luci) gene (Fig 5D). We selected two PbAP2-TR target genes, dihydrolipoamide acyltransferase (*dlat*, PBANKA_0505000) and biotin--protein ligase 1 (*bpl-1*, PBANKA_0511000), for the reporter assays, as these were significantly upregulated upon *pbap2-tr*-disruption (Table S5). The upstream regions of *dlat* and *bpl-1* both contained one of Motif 1 and two of Motif 2 (Fig 5E), and we introduced mutations to alter either or both of these motifs from GTTGY and GTTCT to tTTGY and tTTCT, respectively. These pCen-luc plasmids were then introduced into the wild-type *P. berghei* ANKA strain, and dual luciferase assays were performed at 14 hpi in biological triplicate.

In the assay for the *dlat* promoter, the relative expression of f-luci against R-luci was significantly increased by 23- and 7-fold through mutations into Motif 1 and 2, respectively (*p*-value < 0.01, two-tailed Student’s t-test) (Fig 5F). Furthermore, f-luci/R-luci expression was the highest with mutations into both Motifs 1 and 2 (49-fold higher than that without mutations) (Fig 5F). In the assay for *bpl-1*, a similar result was obtained; promoter activity significantly increased with mutations into either Motif 1 or 2 (by 54- and 5-fold, respectively), and mutations into both motifs induced the highest f-luci/R-luci expression level (83-fold higher than that without mutations) (Fig 5F). Thus, for both reporters, Motifs 1 and 2 both functioned as *cis*-regulatory elements for transcriptional repression, and Motif 1 contributed more to transcriptional repression than Motif 2.

Next, we evaluated whether the functional difference between Motifs 1 and 2 was also observed in the differential expression analysis between *pbap2-tr*-DiCre^Rapa+^ and *pbap2-tr*-DiCre^Rapa-^. We divided the target genes into three groups: targets of a peak with both motifs, only Motif 1, and only Motif 2, within 300 bp of the summit (Table S5). The average log_2_(fold change) value of the “both motifs” group was significantly higher than that of the other two groups with *p*-values less than 0.05 by two-tailed Student’s t-test (Fig 5G). This suggests that transcriptional repression by PbAP2-TR is strong in regions containing both binding motifs. For targets with either motif, the average log_2_(fold change) value was significantly higher in the Motif 1 group than in the Motif 2 group (Fig 5G). These results verify that Motif 1 is a stronger *cis*-regulatory element for transcriptional repression by PbAP2-TR than Motif 2. Collectively, these results suggest that a single transcription factor can generate variations in the transcript levels of target genes through combinations of different *cis*-regulatory elements. In a previous study, we described a similar mechanism for transcriptional activation by PbSIP2 (28). These *cis*-element dependent variations in the levels of transcriptional activation and repression may be important for the parasite to regulate its complex life cycle with a small number of sequence-specific transcription factors (19).

### PbAP2-TR mainly targets ribosome biogenesis-related genes and exported protein genes and contributes toward establishing their transcription peak patterns

PbAP2-TR targets included 458 genes that were functionally annotated in the PlasmoDB (https://plasmodb.org/plasmo/app) (Table S3). To evaluate the role of transcriptional repression by PbAP2-TR in asexual blood stage development, we classified these annotated genes into characteristic groups. Among 13 groups, the “protein biogenesis” group was the largest, containing 94 target genes (Fig 6A). These mainly included ribosome-related genes, such as 40S and 60S ribosomal protein genes and nucleolar protein genes. The target genes also included many genes belonging to “metabolism,” “exported protein,” and “mitochondria” (Fig 6A). We further performed gene ontology (GO) analysis of the PbAP2-TR targets to explore the enriched gene sets. The analysis revealed that the term “ribosome biogenesis” was most enriched in the target genes with a *p*-value of 2.1 × 10^−6^, consistent with the significant number of ribosome-related genes found in the targets (Table S6). Furthermore, several terms related to ribosomes, such as “ribonucleoprotein complex biogenesis,” “rRNA processing,” and “nucleolus,” were included in the 20 most enriched terms (Table S6). Other enriched terms included some related to parasite proteins exported into the host cell (such as “host cellular component” and “symbiont-containing vacuole”) and two IMC-related terms (“pellicle” and “inner membrane pellicle complex”) (Table S6).

**Figure 6.**
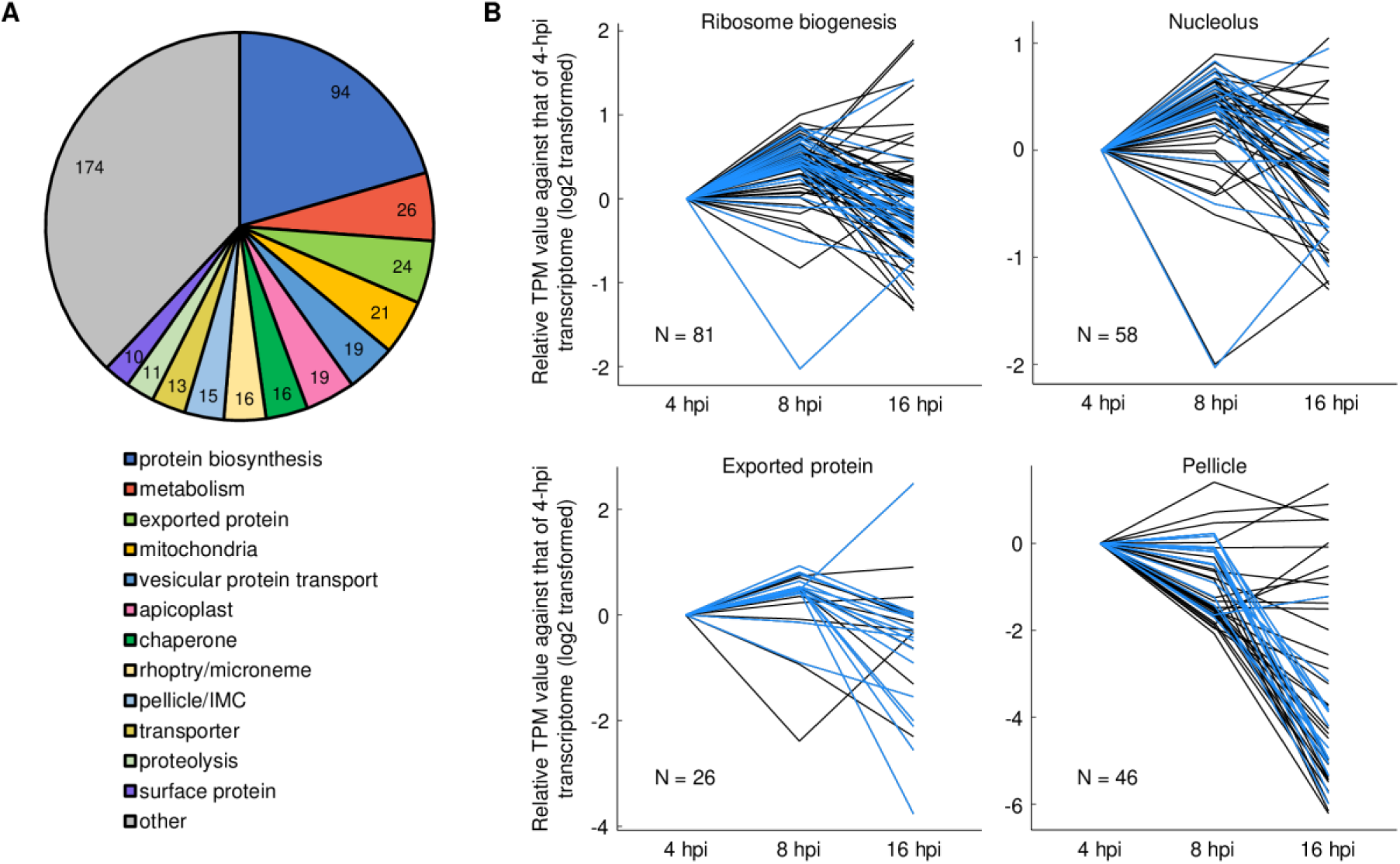
Functional profiling of PbAP2-TR targets and the expression patterns of target-enriched gene sets. (A) Classification of PbAP2-TR target genes into 13 characteristic groups. Genes annotated on the PlasmoDB (https://plasmodb.org/plasmo/app) (458 genes) were classified. (B) Expression patterns of gene sets enriched in the PbAP2-TR targets. Transcripts per million (TPM) values at 4, 8, and 16 hpi are shown as relative values normalized to those of 4 hpi. N indicates the number of genes in each set (genes with TPM less than 30 at all time-points were excluded). The PbAP2-TR targets are highlighted in blue.

In the course of the *Plasmodium* IDC, the expression of most genes changes with cell stage progression rather than being constantly expressed, and genes of the same functional group mostly show similar expression patterns. To investigate whether PbAP2-TR contributes to the establishment of unique expression peak patterns for each functional gene group, we assessed the transcript levels of target-enriched group genes at 4 (ring, before PbAP2-TR expression), 8 (early trophozoite, beginning of PbAP2-TR expression), and 16 hpi (late trophozoite, end of PbAP2-TR expression) (Table S7). For most genes with the GO term “ribosome biogenesis” and “nucleolus,” their transcript per million (TPM) values peaked at 8 hpi when PbAP2-TR expression began (Fig 6B). Similarly, the expression peaks for most *Plasmodium* exported protein genes (including *exp1–3* and *ibis1*) were also observed at 8 hpi (Fig 6B). The “pellicle” genes mostly showed no notable change or a slight decrease from 4 to 8 hpi, but similar to ribosome-related genes, their expression decreased towards 16 hpi (Fig 6B). Collectively, gene groups enriched in PbAP2-TR targets commonly showed patterns of downregulation from 8 to 16 hpi, during which PbAP2-TR is expressed. These results suggest that PbAP2-TR contributes to the establishment of transcription peak patterns in its target genes by repressing their transcription during the IDC.

### PbAP2-TR recruits PbMORC as a co-factor

To further investigate the mechanism of transcriptional repression by PbAP2-TR, we performed rapid immunoprecipitation mass spectrometry (MS) of endogenous proteins (RIME), which involves ChIP, followed by MS analysis, and explored its co-factors. ChIP was conducted using PbAP2-TR::GFP*^pbap2-g^*^(-)^ and WT (as controls) parasites at 12 hpi, as described for the ChIP-seq analyses. IPed proteins were harvested by on-bead digestion with trypsin and subjected to liquid chromatography-tandem MS (LC-MS/MS) analysis. MS data obtained from four biologically independent experiments were compared between PbAP2-TR::GFP*^pbap2-g^*^(-)^ and WT. Through this analysis, we identified five proteins that were unique or more than five-fold enriched in PbAP2-TR::GFP*^pbap2-g^*^(-)^ ChIP compared with those in WT ChIP with *p*-values less than 0.01 by a two-tailed Student’s t-test (Fig 7A and Table S8). Among these, PbAP2-TR itself and PbMORC had average quantitative values higher than 20, whereas the values for the others were lower than 2, suggesting that PbMORC is a strong candidate for a PbAP2-TR co-factor (Fig 7A and Table S8).

**Figure 7.**
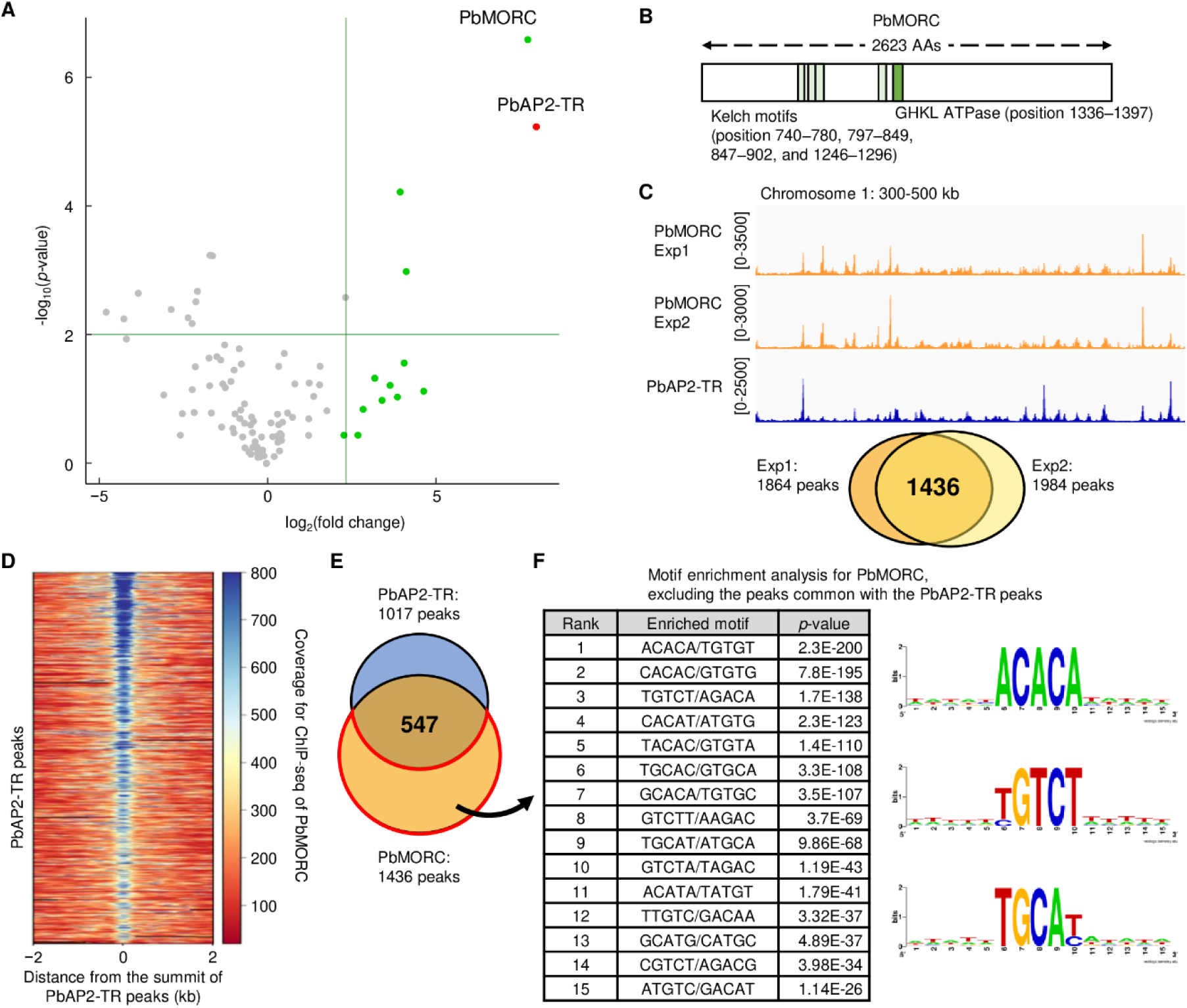
Recruitment of PbMORC by PbAP2-TR as a co-factor. (A) Volcano plot showing proteins enriched in the RIME of PbAP2-TR::GFP compared with that of the wild-type (WT) at 12 hpi. Proteins uniquely identified in the PbAP2-TR::GFP samples are indicated by green dots. To display proteins with an average quantitative value of 0 on the plot, pseudo fold change values were calculated by adding a pseudo-count value (0.1). Horizontal and vertical lines indicate a *p*-value of 0.01 and a fold change value of 5, respectively. (B) Schematic illustration of PbMORC. Regions of the GHKL ATPase and Kelch motif are indicated by green and light green, respectively. (C) Integrative Genomics Viewer (IGV) images for PbMORC ChIP-seq experiments 1 and 2 on chromosome 1. An IGV image of the PbAP2-TR ChIP-seq experiment 1 is shown below. Histograms show the row read coverage of ChIP data at each base. Scales are indicated in square brackets. The Venn diagram at the bottom shows the number of overlapping peaks between experiments 1 and 2. (D) Heat map showing coverage in the ChIP-seq of PbMORC at the PbAP2-TR peaks. Peak regions are aligned in the ascending order of their *q*-values. (E) Venn diagram showing the number of overlapping peaks between PbAP2-TR ChIP-seq and PbMORC ChIP-seq. (F) DNA motifs enriched in PbMORC peak regions that did not overlap with the PbAP2-TR peaks. Logos for the enriched motifs were generated using WebLogo and are displayed on the right.

*Plasmodium* parasites possess a single MORC, which has a conserved GHKL (gyrase, Hsp90, histidine kinase, and MutL)-ATPase domain and several Kelch motifs (Fig 7B) (39). To confirm that PbAP2-TR recruits PbMORC to the upstream regions of target genes as a co-factor, we performed ChIP-seq analysis of PbMORC at 12 hpi. For ChIP-seq, a parasite line expressing GFP-fused PbMORC was generated, and *pbap2-g* was disrupted in this parasite [PbMORC::GFP*^pbap2-g^*^(-)^, Fig S4] to compare the ChIP-seq data with that of PbAP2-TR::GFP*^pbap2- g^*^(-)^. Duplicate experiments identified 1864 and 1984 peaks, of which 1436 peaks (77% of the experiment 1 peaks) overlapped (Fig 7C, Table S9A and S9B). The ChIP-seq coverage of PbMORC showed peaks at the summit of most PbAP2-TR peaks, indicating genome-wide co-localization of the two factors in the PbAP2-TR binding regions (Fig 7D). In addition, Motifs 1 and 2 were significantly enriched in the PbMORC peak regions with *p*-values of 1.5 × 10^−52^ and 0.035, respectively, by Fisher’s exact test (Table S9C). These results indicate that PbAP2-TR recruits PbMORC to the upstream regions of its target genes. Importantly, while ChIP-seq showed co-localization in the PbAP2-TR binding regions, only 38% of the PbMORC peaks (547 peaks) overlapped with the PbAP2-TR peaks (Fig 7E). Motif enrichment analysis of the remaining PbMORC peaks (889 peaks) detected ACACA, TGCAY, and YGTCT, along with their related motifs (Fig 7F and Table S9D). This suggests that transcription factors recognizing these motifs may also recruit PbMORC as a co-factor.

## Discussion

In *Plasmodium*, transcriptional repressors have been shown to play a role in regulating cell fate bifurcations, namely sexual differentiation and sex determination, and in silencing subtelomeric multigene family expression (22, 27, 30, 40–43). These processes are regulated in a mutually exclusive or clonally variant manner and often involve trimethylated histone H3 Lys9, which is a hallmark of heterochromatin (44–46). In contrast, asexual blood stage development is a simple replication cycle that proceeds without cell fate diversion. Thus, it could be considered that transcriptional regulation during the IDC does not require transcriptional repressors and proceeds via a cascade of transcriptional activators alone that induce each stage-specific transcription in a “just-in-time” manner. However, in this study, we demonstrated that the transcriptional repressor PbAP2-TR is essential for the progression of asexual blood stage development. The target genes of PbAP2-TR include many genes related to protein biosynthesis, mainly ribosome biogenesis, and exported protein genes. Ribosome biogenesis is the first step in protein biosynthesis (47), and exported proteins are expressed to modify host RBCs, which is an important first step for intraerythrocytic development (48, 49). The transcription pattern of these genes peaks during the ring to trophozoite development and decreases towards the late trophozoite stage as their requirement for the later cell stage decreases (50, 51). Our results indicate that PbAP2-TR establishes this transcription peak pattern by repressing their transcription in the course of early to late trophozoite development, thereby being essential for promoting transition from the cell growth to replication phases of the IDC. This role of PbAP2-TR suggests that the “just-in-time” gene expression patterns during IDC are strictly controlled at the transcriptional level through a combination of transcriptional activators and repressors, providing an insight into the role of transcriptional repressors in asexual blood stage development.

In the PbAP2-TR targets, we also detected enrichment of pellicle/IMC-related genes. The pellicle and IMC structures are important for merozoite formation and motility (52–54), and genes related to these structures are included among the targets of PbSIP2, a master transcription factor that regulates merozoite formation (28). Consistently, their expression in the IDC reaches a peak during schizont development and is downregulated after RBC invasion (14). Given this, the role of PbAP2-TR is not limited to the establishment of transcription peak patterns through repressing genes that are highly transcribed during early trophozoite development; it also represses genes that are not required for or are possibly harmful to trophozoite development. In a previous study, we revealed that the female transcriptional repressor PbAP2-FG2 plays multiple roles in promoting female development, such as repression of early gametocyte genes and suppression of male fate (27). These suggest that in *Plasmodium*, transcriptional repressors target wide-variety of genes to play multiple roles, unlike transcriptional activators, whose function is limited to regulate certain stage-specific genes.

MORC family proteins are conserved in eukaryotes, including apicomplexan parasites, and play roles in gene silencing and chromatin compaction (55). *Plasmodium* parasites have a single MORC ortholog that is essential for asexual blood stage development and is proposed to be involved in chromatin organization and heterochromatin-related gene silencing (39, 56). Besides this broad-scale chromatin regulation, the role of MORC has also been suggested as a transcriptional regulator associated with AP2 transcription factors; *i.e.*, MORC was also identified through co-IP with some AP2 transcription factors, and co-IP with PfMORC detected several other AP2 transcription factors (27, 30, 39). Such interactions between the MORC protein and AP2 transcription factors have also been detected in other apicomplexan parasite, *Toxoplasma* (57–61). In this study, we performed a ChIP-seq analysis of PbMORC and demonstrated that PbMORC is genome-widely recruited to a specific *cis*-regulatory element by PbAP2-TR. These results strongly indicate that MORC is involved as a co-factor in the transcriptional regulation of genes downstream of transcription factor binding sites. In *Toxoplasma*, MORC functions in a complex with histone deacetylase 3 to regulate sexual commitment, suggesting that MORC functions are related to histone modifications (57). In contrast, RIME analysis with PbAP2-TR detected PbMORC as a major co-factor, but no histone-modifying enzymes. Similarly, our previous study revealed PbMORC as the only major co-factor of the female-specific transcriptional repressor complex, PbAP2-FG2/PbAP2R-2 (27). These indicate that in *Plasmodium*, MORC does not require other co-factors for its functions but represses target gene expression simply via remodeling chromatin states as a possible chromatin remodeler.

In summary, our results indicate that transcriptional repression is essential for establishing precise transcriptional profiles during *Plasmodium* asexual blood stage development. In the IDC transcriptomes, several gene groups show different peak patterns of gene transcription, which constitutes the cascade of “just-in-time” transcription, suggesting that multiple transcriptional repressors are involved in generation of this cascade. *Plasmodium* parasites have several AP2 transcription factors essential for asexual blood stage development (32, 34), some of which have not been functionally investigated. As our ChIP-seq of PbMORC detected the enrichment of several motifs other than PbAP2-TR-binding motifs, these factors may include other transcriptional repressors that recruit PbMORC. We believe that further exploration of their functions is important for understanding the processes of parasitic blood stage development.

## Materials and Methods

### Ethics statement

All experiments were performed in accordance with the recommendations of the Guide for the Care and Use of Laboratory Animals of the National Institutes of Health to minimize animal suffering and were approved by the Animal Research Ethics Committee of Mie University (permit number 23–29).

### Parasite preparation

The parasites were inoculated into ddY mice. All parasites used in this study were derived from the wild-type (WT) *P. berghei* ANKA strain. Transgenic parasites generated using the CRISPR/Cas9 method were derived from the WT-originated Cas9-expressing parasite, PbCas9, which has a Cas9 expression cassette at the *p230p* locus (27). The *pbap2-tr*-DiCre parasite was generated from PbCas9^DiCre^, which constitutively expresses Cre59 (Thr19–Asn59) fused with FKBP12 and Cre60 (Asn60–Asp343) fused with FRB in addition to Cas9 (28).

*In vitro* cultures of whole blood from infected mice were performed using RPMI1640 medium supplemented with 25% fetal calf serum and penicillin/streptomycin at 37 °C in 5% CO_2_ and 10% O_2_. For cell cycle synchronization, infected blood was cultured for 16 h, and mature schizonts produced in the cultures were harvested by density gradient centrifugation using an iodixanol solution (Optiprep), whose density was adjusted to 1.077 g/mL by mixing with tricine solution. The harvested schizonts were then intravenously injected into mice. Parasitemia and parasite morphology were assessed on Giemsa-stained blood smears.

### Generation of transgenic parasites via the CRISPR/Cas9 system

For gene editing with the CRISPR/Cas9 system using PbCas9 and PbCas9^DiCre^, donor DNAs and single guide RNA (sgRNA) vectors were constructed as previously described (35). Briefly, templates for donor DNAs were constructed using overlap PCR and cloned into pBluescript KS (+) by In-Fusion cloning (Takara) at the *Xho*I and *Bam*HI sites. Donor DNAs were then amplified by PCR from the constructed plasmid. The target sequences of the sgRNAs were designed using CHOPCHOP (https://chopchop.cbu.uib.no/). sgRNA templates were constructed by annealing DNA oligos and were cloned into the sgRNA vector using the DNA Ligation Kit (Takara).

Transfection was performed using cultured schizonts and DNA constructs described above. Briefly, recipient parasites were cultured for 16 h, and schizonts were harvested by density gradient centrifugation. The harvested schizonts were then transfected with DNA constructs using the Amaxa Basic Parasite Nucleofector Kit 2 (LONZA) and were immediately inoculated into mice. All transfectants were selected by treating the infected mice with 70 μg/mL pyrimethamine in their drinking water. For CRISPR/Cas9 gene editing, recombination was confirmed using PCR and/or Sanger sequencing, and clonal parasites were obtained by limiting dilution. All the primers used in this study are listed in Table S10.

### Fluorescence microscopic analysis

Fluorescence analysis was performed using the Olympus BX51 microscope with Olympus DP74 camera. For nucleus detection, parasites were stained with 1 ng/mL Hoechst 33342 for 10 min at 37 °C.

### Conditional knockout of *pbap2-tr*

Whole blood from mice infected with *pbap2-tr*-DiCre (or *pbap2-tr*-DiCre*^pbap2-g^*^(-)^) parasites was cultured. The cultures were then split into two parts; a 3/20 volume of rapamycin (Wako) solution (200 μM DMSO stock) was added to one part (final concentration of 30 nM), and the same volume of DMSO was added to the other as a control. The parasites were cultured for 16 h, and schizonts were harvested and inoculated into mice. For *in vitro* phenotype analysis, the parasites were cultured again at 10 hpi. For *in vivo* phenotype analysis, parasites were grown in mice for 3 days, and parasitemia was assessed daily by Giemsa staining. Genotyping PCR to confirm recombination at the *pbap2-tr* locus was performed at 16 h after starting the cultures before schizont harvest and on day 3 of the *in vivo* phenotype analysis. The primers used for genotyping are listed in Table S10.

### ChIP-seq and sequencing data analysis

ChIP-seq was performed as previously described (28). Briefly, infected blood was passed through a Plasmodipur filter, and formalin solution was immediately added to a final concentration of 1% for cell fixation. After 1 h of fixing at 30 °C, RBCs were lysed in ice-cold 1.5 M NH_4_Cl solution, and residual cells were lysed in SDS lysis buffer. The cell lysate was sonicated using the Bioruptor (Cosmo Bio), and immunoprecipitation was performed using anti-GFP polyclonal antibodies (Abcam) conjugated to Protein A Magnetic Beads (Invitrogen). Libraries were constructed from DNA fragments purified from the immunoprecipitated chromatin using the KAPA HyperPrep Kit (Kapa Biosystems). Next-generation sequencing (NGS) was performed using the MGI DNBSEQ-G400. DNA fragments purified from cell lysates before immunoprecipitation were also sequenced as input sequence data. Two biologically independent experiments were performed for each ChIP-seq experiment.

Reads in the sequence data were mapped onto the reference genome sequence of *P. berghei* (v3.0, downloaded from PlasmoDB) using Bowtie 2. Those aligned onto more than two sites in the genome were removed from the mapping data, and peaks were called by the macs2 callpeak using the input sequence data as control (fold enrichment > 3.0, *q*-value < 0.01, and setting --call-summit option on). The parameters for all programs were set as default unless otherwise indicated. Common peaks in duplicates were defined as those with a distance of less than 150 bp between their peak summits.

The enrichment of motifs within 50 bp of the peak summits was analyzed using Fisher’s exact test. For this motif enrichment analysis, peaks located within 1 kb of the end of each chromosome were excluded to avoid detection of the telomeric repeat sequence TTYAGGG (Y = T or C). In the *P. berghei* reference genome, 13 of 28 chromosome end sequences are fully resolved, and these include less than 1-kb telomeric repeat sequences. Repeats of TTYAGGG are also found within subtelomeric 2.3-kb repeat units. Genes with peaks within 1200 bp of their start codons were identified as target genes.

### DIP-seq analysis

DIP-seq was performed as previously described (25). Briefly, the sequences for the AP2 domains of PbAP2-TR (164–336 for AP2 domains 1–2 and 1458–1527 for AP2 domain 3) were cloned into the MBP fusion vector pMal-c5X (NEB). *E. coli* transformed with the plasmids was cultured for 12 h at 37 °C, and the expression of MBP-fused proteins was then induced with isopropyl β-D-thiogalactopyranoside (final concentration of 200 nM). Recombinant AP2 domains fused with MBP were purified using amylose resin (NEB) and were mixed with *P. berghei* ANKA genomic DNA fragments. After 30 min of incubation at room temperature, the recombinant proteins and bound DNA fragments were purified using amylose resin. DNA fragments were then subjected to library preparation and NGS using the Illumina NextSeq 500, as described for the ChIP-seq analysis. Genomic DNA fragments were sequenced as inputs before use in the DIP. Sequence data were analyzed similar to that in ChIP-seq, except for the parameters of peak calling (fold enrichment > 2.5, *q*-value < 0.01). Motif enrichment analyses were performed separately for three chromosome sets (chromosomes 1–8, 9–12, and 13–14) because the analysis using all DIP-seq peaks detected several motifs with *p*-values less than 5.0 × 10^−324^, which is the smallest positive real number on the R platform.

### RNA-seq and sequence data analysis

Rapamycin-dependent disruption of *pbap2-tr* was induced, and schizonts of *pbap2-tr*-DiCre^Rapa-^ and *pbap2-tr*-DiCre^Rapa+^ were inoculated into mice, as described for the conditional knockout of *pbap2-tr*. At 12 hpi, whole blood infected with *pbap2-tr*-DiCre^Rapa-^ or *pbap2-tr*-DiCre^Rapa+^ parasites was passed through a Plasmodipur filter. RBCs were lysed in ice-cold 1.5 M NH_4_Cl solution, and total RNA was extracted from the residual parasites using the Isogen II reagent (Nippon Gene). RNA-seq libraries were prepared using the KAPA mRNA HyperPrep Kit (Kapa Biosystems) and were sequenced using the MGI DNBSEQ-G400. Three biologically independent experiments were performed for *pbap2-tr*-DiCre^Rapa-^ and *pbap2-tr*-DiCre^Rapa+^. The sequence data were mapped onto the reference *P. berghei* ANKA genome sequence using HISAT2, with the maximum intron length set as 1000. The number of reads mapped to each gene was calculated using featureCounts and compared between *pbap2-tr*-DiCre^Rapa-^ and *pbap2-tr*-DiCre^Rapa+^ using DESeq2. Genes with an average TPM < 30 in both *pbap2-tr*-DiCre^Rapa-^ and *pbap2-tr*-DiCre^Rapa+^ and subtelomeric multigene families (*pir* and *fam*) were removed from the differential expression analysis. Default parameters were set for all programs unless indicated otherwise.

### Dual-luciferase reporter assay

Plasmids were constructed to generate luciferase reporter lines as previously described (22). Briefly, the upstream regions of *dlat* and *bpl-1* were amplified from *P. berghei* genomic DNA by PCR and inserted upstream of the firefly luciferase gene in the reporter plasmid pCen-luc using the *Kpn*I and *Nhe*I sites. In addition to the firefly luciferase gene, pCen-luc contains the Renilla luciferase gene expression cassette as an internal control and the human *dhfr* gene as a drug-selectable marker. The expression of these two genes is under the control of the bidirectional *P. berghei* elongation factor 1-alpha gene promoter. The plasmid also contains the centromere sequence from the *P. berghei* chromosome 5 (62). Mutations within *dlat* and *bpl-1* promoters were introduced using overlap PCR before integration of their upstream sequences into the reporter plasmid.

Parasites carrying pCen-luc were synchronized as described in the parasite preparation method and were harvested from the tail vein of infected mice at 14 hpi. RBCs were lysed in ice-cold lysis solution (1.5 M NH_4_Cl, 10 mM EDTA, and 0.1 M KHCO_3_) for 5 min, and the residual cells were subjected to dual-luciferase reporter assays using the Dual-Luciferase Reporter Assay System (Promega). The assays were performed according to the manufacturer’s instructions. Firefly and *Renilla* luciferase expression was assessed using the GloMax 96 Microplate Luminometer (Promega). The assays were performed in biological triplicate for each sample.

### RIME

RIME was performed as previously described (27). Briefly, ChIP was performed as described for ChIP-seq analysis using PbAP2-TR::GFP, and protein complexes bound on Protein A beads were digested overnight at 37 °C with trypsin (Promega) and further incubated for 4 h at 37°C with additional trypsin. The peptides released from the beads through digestion were then purified using a C18 tip (GL-Science, Tokyo, Japan) and then subjected to nanocapillary reversed-phase LC-MS/MS analysis using a C18 column (12 cm × 75 μm, 1.9 μm, Nikkyo Technos, Tokyo, Japan) on a nanoLC system (Bruker Daltoniks, Bremen, Germany) connected to a timsTOF Pro mass spectrometer (Bruker Daltoniks) and a modified nano-electrospray ion source (CaptiveSpray; Bruker Daltoniks). From the MS/MS data, IPed proteins were identified using DataAnalysis version 5.2 (Bruker Daltoniks) and MASCOT version 2.7.0 (Matrix Science, London, UK) against the Uniprot_Plasmodium_berghei_ANKA_strain database (4948 sequences; 3412795 residues). Protease specificity was set for trypsin (C-term, KR; Restrict, P; Independent, no; Semispecific, no; two missed and/or nonspecific cleavages permitted); conversion of the N-terminal Gln to pyro-Glu and oxidation of methionine were considered as possible modifications. The mass tolerance for precursor ions and fragment ions were ±15 ppm and ±0.05 Da, respectively. The threshold score/expectation value for accepting individual spectra was P < 0.05. Quantitative values were determined using Scaffold5 version 5.1.2 (Proteome Software, Portland, OR, USA). Four biologically independent experiments were performed for PbAP2-TR::GFP and the WT. Fold change and *p*-value (two-tailed Student’s t-tests) were calculated using Microsoft Excel.

## Supplementary Data

**Fig. S1.**
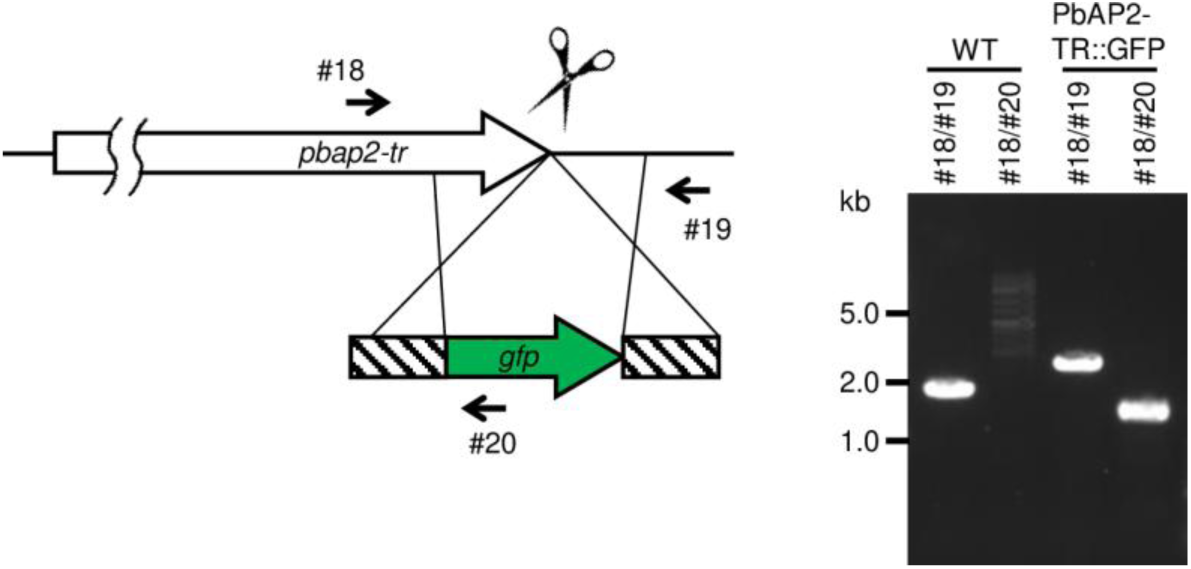
Genotyping of PbAP2-TR::GFP. Schematic illustration of gene editing at the *pbap2-tr* locus is shown on left. A gel image from the genotyping PCR analysis is shown on right. The primer numbers are listed in Table S8.

**Fig. S2.**
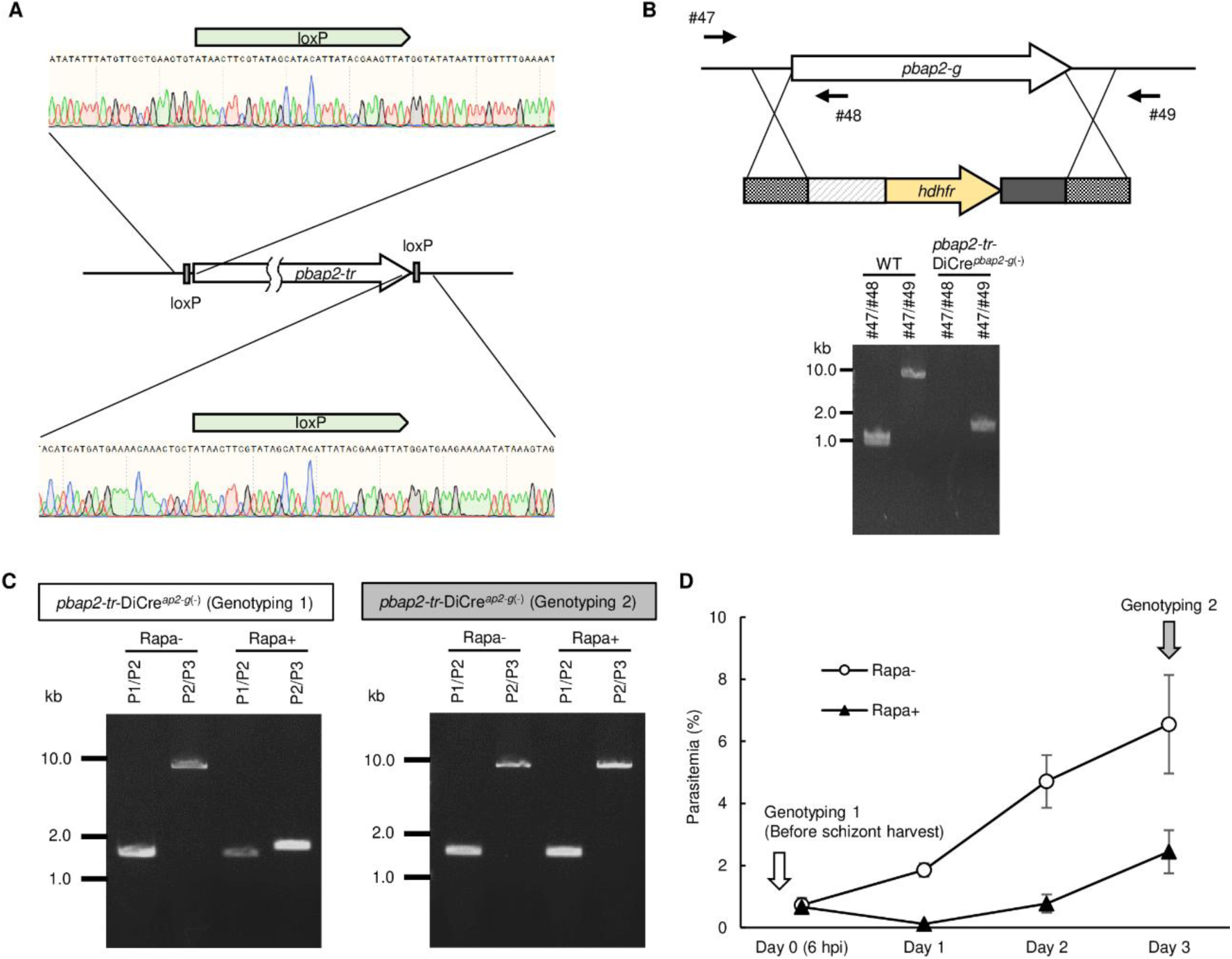
Generation of of *pbap2-tr*-DiCre and *pbap2-tr*-DiCre*^ap2-g^*^(-)^. (A) Genotyping of *pbap2-tr*-DiCre. Sangar sequence results confirming insertion of loxP at 5′- and 3′-side of *pbap2-tr* are shown on top and bottom. A schematic illustration of the *pbap2-tr* locus in *pbap2-tr*-DiCre is shown between them. (B) Genotyping of *pbap2-tr*-DiCre*^pbap2-g^*^(-)^. Schematic illustration of gene editing at the *pbap2-g* locus is shown on top. A gel image from the genotyping PCR analysis is shown on bottom. The primer numbers are listed in Table S8. (C) Representative gel images of the genotyping PCR analysis for *pbap2-tr*-DiCre*^ap2-g^*^(-)_Rapa−^ and *pbap2-tr*-DiCre*^ap2-g^*^(-)_Rapa+^ performed at 16 h after starting culture (Genotyping 1, left) and on Day 3 after inoculating them into mice (Genotyping 2, right). Primers used are illustrated in Fig 2A. (D) Parasite growth of *pbap2-tr*-DiCre*^ap2-g^*^(-)_Rapa−^ and *pbap2-tr*-DiCre*^ap2-g^*^(-)_Rapa+^ *in vivo*. Parasitemia was assessed by Giemsa staining. Error bars indicate the standard error of mean value from three biologically independent experiments. Time points for Genotyping 1 and 2 are indicated by arrows.

**Fig. S3.**
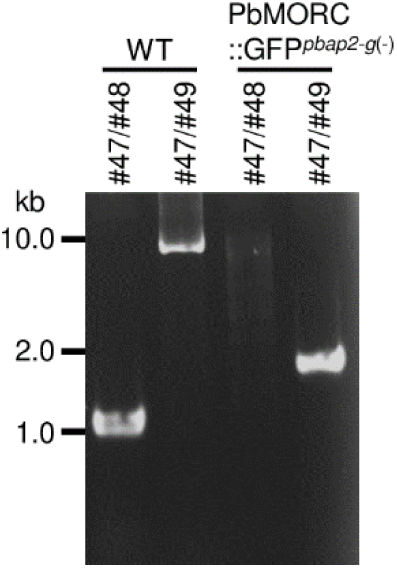
Genotyping of PbAP2-TR::GFP*^pbap2-g^*^(-)^. PbAP2-TR::GFP*^pbap2-g^*^(-)^ was derived from PbAP2-TR::GFP. *pbap2-g* was disrupted by the same strategy illustrated in Fig S2B. The primer numbers are listed in Table S8.

**Fig. S4.**
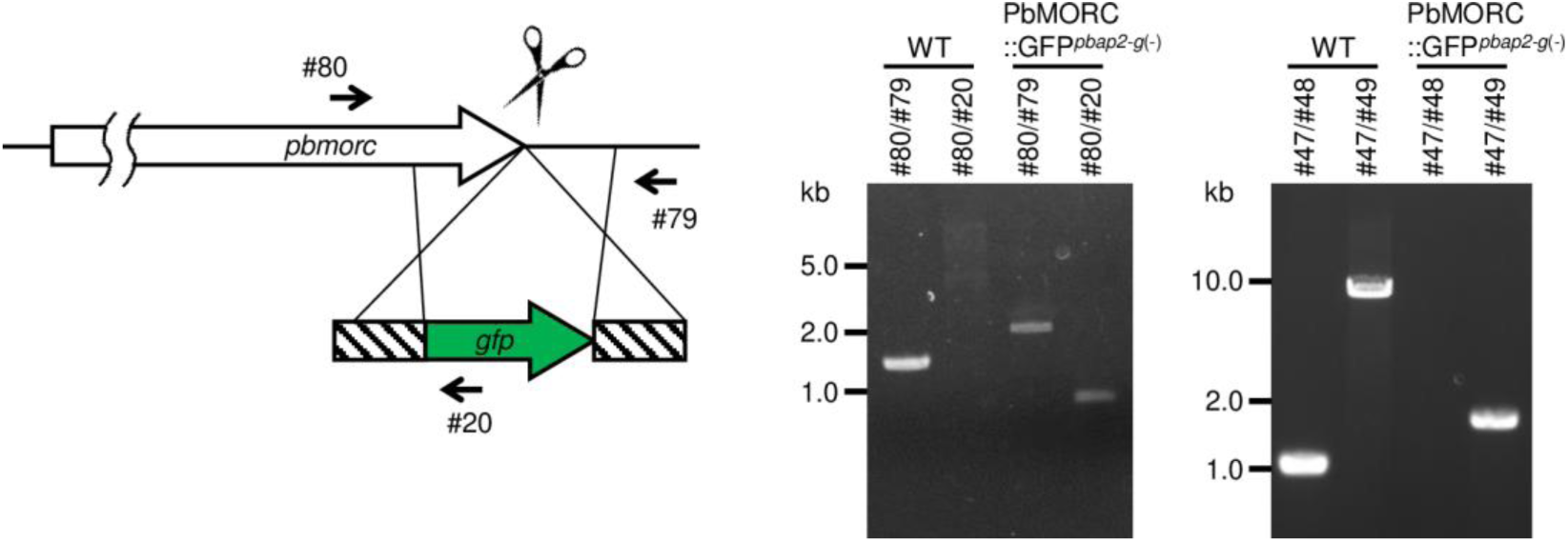
Genotyping of PbMORC::GFP*^pbap2-g^*^(-)^. Schematic illustration of gene editing at the *pbmorc* locus is shown on left. Gel images from the genotyping PCR analysis for *pbmorc* and *pbap2-g* loci are shown on right. *pbap2-g* was disrupted by the same strategy illustrated in Fig S2B. The primer numbers are listed in Table S8.

**Table S1. ChIP-seq analysis of PbAP2-TR**

(A) Peaks in experiment 1. (B) Peaks in experiment 2. (C) Motif enrichment analysis.

**Table S2. DIP-seq analysis of PbAP2-TR**

(A) DIP-seq peaks for AP2 domain 1–2. (B) Motif enrichment analysis for DIP-seq of AP2 domain 1–2. (C) DIP-seq peaks for AP2 domain 3. (D) Motif enrichment analysis for DIP-seq of AP2 domain 3.

**Table S3. Target genes of PbAP2-TR**

**Table S4. Differential expression analysis between 8 and 16 hpi using *ap2-g*-knockout parasite**

**Table S5. Differential expression analysis between *pbap2-tr*-DiCre^Rapa-^ and *pbap2-tr*-DiCre^Rapa+^**

**Table S6. Gene ontology analysis of PbAP2-TR targets**

**Table S7 Time-course transcriptomic analysis using *ap2-g*-knockout parasite**

**Table S8. RIME analysis using PbAP2-TR::GFP and WT parasites**

**Table S9. ChIP-seq analysis of PbMORC**

(A) Peaks in experiment 1. (B) Peaks in experiment 2. (C) Motif enrichment analysis with all ChIP-seq peaks. (D) Motif enrichment analysis excluding peaks common with PbAP2-TR peaks.

**Table S10. List of primers used in this study**

